# Physical activity and aerobic fitness show different associations with brain processes underlying anticipatory selective visuospatial attention in adolescents

**DOI:** 10.1101/2020.07.29.227181

**Authors:** Doris Hernández, Erkka Heinilä, Joona Muotka, Ilona Ruotsalainen, Hanna-Maija Lapinkero, Heidi Syväoja, Tuija H. Tammelin, Tiina Parviainen

**Affiliations:** Department of Psychology, Center for Interdisciplinary Brain Research, University of Jyväskylä, Mattilanniemi 6, FI-40014, Jyväskylä, Finland; LIKES Research Center for Physical Activity and Health, Jyväskylä, Rautpohjankatu 8, FIN-40700 Jyväskylä Finland; Department of Psychology, University of Jyväskylä, Mattilanniemi 6, FI-40014, Jyväskylä, Finland

**Keywords:** physical activity, aerobic fitness, adolescence, anticipatory alpha oscillations, selective attention, magnetoencephalography

## Abstract

Underlying brain processes of exercise-related benefits on executive functions and the specific contribution of physical activity vs. aerobic fitness are poorly understood, especially during adolescence. We explored whether and how physical activity and aerobic fitness are associated with selective attention and the oscillatory dynamics induced by an anticipatory spatial cue. Further, we studied whether the link between physical exercise level and cognitive control in adolescents is mediated by the task-related oscillatory activity. Magnetoencephalographic alpha oscillations during a modified Posner’s cueing paradigm were measured in 59 adolescents (37 females and 22 males, 12 to 17 years). Accelerometer-measured physical activity and aerobic fitness (20-m shuttle run test) were used to divide the sample into higher and lower performing groups. The interhemispheric alpha asymmetry during selective attention was larger in the high than in the low physical activity group, but there was no difference between the high and low aerobic fitness groups. Exploratory mediation analysis suggested that anticipatory interhemispheric asymmetry mediates the association between physical activity status and drift rate in the selective attention task. Higher physical activity was related to increased cue-induced asymmetry, which in turn was associated with less efficient processing of information. Behaviorally, higher physically active males showed stronger dependence on the cue while higher fit females showed more efficient processing of information. Our findings suggest that physical activity may be associated with a neural marker of anticipatory attention in adolescents. These findings have implications for understanding the varying results on the association between physical activity and attention in adolescents.

**HIGHLIGHTS:**

- Physical activity and aerobic fitness link differently with attention in adolescents.
- Higher physical activity relates with stronger cue-induced oscillatory asymmetry.
- Brain hemispheric interaction mediates the link between physical activity and attention.

## INTRODUCTION

Recent research has suggested a positive influence of higher levels of physical activity (PA) and aerobic fitness (AF) on cognitive functions (Hillman et al., 2008; Smith et al., 2010). Research has focused so far in the early and late years of life, in pre-adolescent children (Crova et al., 2014; Haapala et al., 2014; Syväoja et al., 2013; van der Niet et al., 2014) and older adults (Colcombe and Kramer, 2003). In children, higher levels of PA and AF have been linked with better academic performance (Haapala et al., 2014; Syväoja et al., 2013), as well as different aspects of cognition/executive functions (Crova et al., 2014; Syväoja et al., 2014; van der Niet et al., 2014). Especially, attention (Syväoja et al., 2014) and inhibitory control (Crova et al., 2014; van der Niet et al., 2014) seem to be associated with increased PA or AF levels in children.

Neural mechanisms underlying the possible advantage in attentional/inhibitory processes because of regular PA and higher AF are still unknown. Previous studies at the microscopic (molecular and cellular) and mesoscopic (circuit) levels, mostly done in mice, report changes in the hippocampus associated with regular exercise (for a review, see Wang and van Praag, 2012). PA is considered an important metabolic activator significantly increasing the mRNA levels of some neurotrophic factors such as the brain-derived neurotrophic factor (BDNF) (Gomez-Pinilla et al., 2008) or the vascular-endothelial growth factor (Carro et al., 2001; Fabel et al., 2003). These growth factors lead to structural and functional changes in the brain, such as neurogenesis (Brown et al., 2003), enhanced synaptic plasticity (O’Callaghan et al., 2007; van Praag et al., 1999), enhanced spine density (Redila and Christie, 2006; Stranahan et al., 2007), and angiogenesis (Carro et al., 2001; van Praag et al., 2005). Indeed, it has also been suggested that PA and AF are linked with these structural changes in the human brain (for a review see Chaddock et al., 2011).

To date, the studies about the effects of either PA or AF over cognition, especially in adolescence (ranging between 10 and 19 years according to the World Health Organization (WHO, 2018), are rarer (Huang et al., 2015; Marchetti et al., 2015; Stroth et al., 2009; Tee et al., 2018; Vanhelst et al., 2016; Westfall et al., 2018). Importantly, the beneficial influence of higher AF (Huang et al., 2015; Hogan et al., 2013, 2015; Marchetti et al., 2015; Stroth et al., 2009; Westfall et al., 2018) or PA (Booth et al., 2013; Tee et al., 2018; Vanhelst et al., 2016) on cognition during adolescence implicates multiple factors. There is some evidence that biological sex modulates the association between PA or AF and cognition in older adult populations (Barha et al., 2017; Baker et al., 2010; Colcombe and Kramer, 2003; van Uffelen et al., 2008). Indeed, Colcombe and Kramer’s (2003) review on the enhancing effects of AF in older adults found that studies including more than 50% of women showed greater cognitive benefits from AF than studies enrolling mostly men. These sex-related differences may influence the link between PA or AF and cognition in other stages of life. Interestingly, this effect could be especially pronounced during adolescence, since this has been suggested to be a sensitive period for pubertal hormone-dependent brain organization (Sisk and Zehr, 2005).

To better understand the effect of PA- and AF-related changes in body metabolism on brain functioning, it is important to consider the specific characteristics of these measures. Even though PA and AF are closely related, they are different concepts and are measured distinctively. PA is related to energy expenditure resulting from any body movement (Caspersen et al., 1985) and it is usually measured by quantifying the amount of movement done in a certain period of time. On the other hand, AF is conceived as an achieved condition and it is usually directly measured by obtaining the maximal oxygen uptake (VO_2_max) during maximal effort tasks. Very few studies have investigated the effect of both PA and AF on cognition in humans in the same study (but see Haapala et al., 2014; Iuliano et al., 2015; Ruotsalainen et al., 2020), especially in adolescents (Tee et al., 2018). A recent study using structural MRI reported the associations of AF, but not accelerometer-measured PA, with grey matter volume in the adolescent’s brain (Ruotsalainen et al., 2019), indicating that the level of PA alone is not directly associated with structural modifications in the brain. Indeed, PA may influence cognition by modulating the dynamics of functional connections rather than structural properties or connectivity in the brain. Although this modulation is difficult to measure, the millisecond temporal resolution of time-sensitive neuroimaging techniques and the analysis of neural oscillations in the brain offer a good way forward for their assessment. While structural measures seem to reflect AF-related effects, it may well be that functional measures, provided by electroencephalography (EEG) and magnetoencephalography (MEG), would be more sensitive to indicate PA-related associations in the brain. Investigating the distinctive effects of PA and AF on cognition and brain function in adolescents could allow a better understanding of the more detailed relationship between body physiology, metabolism and brain function in youth.

Among different cognitive functions, PA and AF have been most often associated with improvements in attention and/or inhibition (Booth et al., 2013; Huang et al., 2015; Stroth et al., 2009; Vanhelst et al., 2016; Westfall et al., 2018). To confirm whether the link between PA or AF and attention/inhibitory control in adolescents reflects underlying modifications in brain function, neuroimaging tasks, specifically tapping on these cognitive functions, need to be applied. By focusing on oscillatory activity, MEG can be used to follow the allocation of attention and the closely linked ability to inhibit irrelevant information. Brain oscillations refer to the rhythmic fluctuations that, at the neural level, represent cyclic changes in the excitability of neuronal populations (Thut et al., 2012) and are suggested to play a key role in coordinating neuronal ensembles necessary for cognitive processing (Lakatos et al., 2008). A widely studied brain oscillation, the alpha rhythm (at around 10 Hz), has been suggested especially important for directing neuronal resources during tasks that require attending or inhibiting some incoming information (Jensen and Mazaheri, 2010; Jensen et al., 2002; Bonnefond and Jensen, 2013; Klimesch, 1999, 2012; Klimesch et al., 2007). It is suggested that alpha oscillations grant access to process relevant information or prevent handling irrelevant information. Thus, the increase of alpha activity is considered as reflecting system-level inhibition in the brain while the suppression of alpha activity is considered as a release from inhibition or the allocation of attention to relevant aspects of a task (Klimesch, 2012). This phenomenon has been successfully used to point to neural mechanisms underlying anticipatory visual attention in general (Jensen and Mazaheri, 2010; Vollebregt et al., 2015) and to changed attentional capacity in attention deficit disorder (Vollebregt et al., 2016). This visuospatial covert attention paradigm may offer a sensitive enough measure to demonstrate changes in attentional resources related to variance in the level of PA or AF. MEG, and especially alpha oscillations, have also shown sensitivity to identifying individual variation (Haegens et al., 2014). To our knowledge, no previous studies have utilized MEG in investigating the associations between PA or AF and anticipatory attention allocation/inhibition processes in the brain of typically developing adolescents.

Although attention/inhibition processes are reported to be influenced by PA or AF, there are clear controversies surrounding the specific behavioral benefits gleaned from PA or AF. Some studies report an increase in accuracy scores (Hogan et al., 2013), while others have found reaction time (RT) reductions (Huang et al., 2015) or both in adolescents (Westfall et al., 2018). Westfall et al. (2018) have suggested a new analysis approach (the EZ diffusion model) to better qualify these measures. In addition to typical task performance measures (accuracy and RT), the EZ model (Wagenmakers et al., 2007) combines accuracy, mean RT and RT variance outcomes to measure dissociable underlying processes used to solve tasks. More specifically, “drift rate” has been defined as an index for the signal to noise ratio of the information processing system to quantify the subject’s ability or task difficulty (Wagenmakers et al., 2007). Further, “boundary separation” is considered a mechanism of speed– accuracy trade-off and response strategy. Finally, “non-decision time” refers to the time period before the decision, which is usually assumed to be included in RT (Luce, 1986). Improvements in drift rate and boundary separation have earlier been related to higher AF in adolescents (Westfall et al., 2018), but the associated brain processes involved in their improvement have not been elucidated.

Here, we explored the link between PA or AF and the neuromagnetic indicators of anticipatory attention/inhibitory control in typically developing adolescents. We recorded alpha oscillations using MEG during a visuospatial selective attention task where subjects had to detect a target (from two stimuli located in the right or left hemifields) anticipated by a preceding cue. We focused on the cue-induced modulation of alpha power in ipsilateral and contralateral hemispheres. Behavioral performance in attention and inhibitory control was also tested. We hypothesized that higher levels of PA or AF would show stronger modulation of the alpha band reflecting more efficient attentional and inhibitory processes in the brain. We thus also tested the exploratory hypothesis that anticipatory alpha oscillations mediate the link between PA or AF and behavioral scores of attention/inhibitory control.

## RESULTS

### Demographic information, physical activity and aerobic fitness

Table 1 summarizes the demographic information for the two groups based on PA and AF. Demographic information is also given separately to males and females. Statistically significant differences between PA groups and sexes (independent-samples t-tests) are denoted.

**Table 1.**
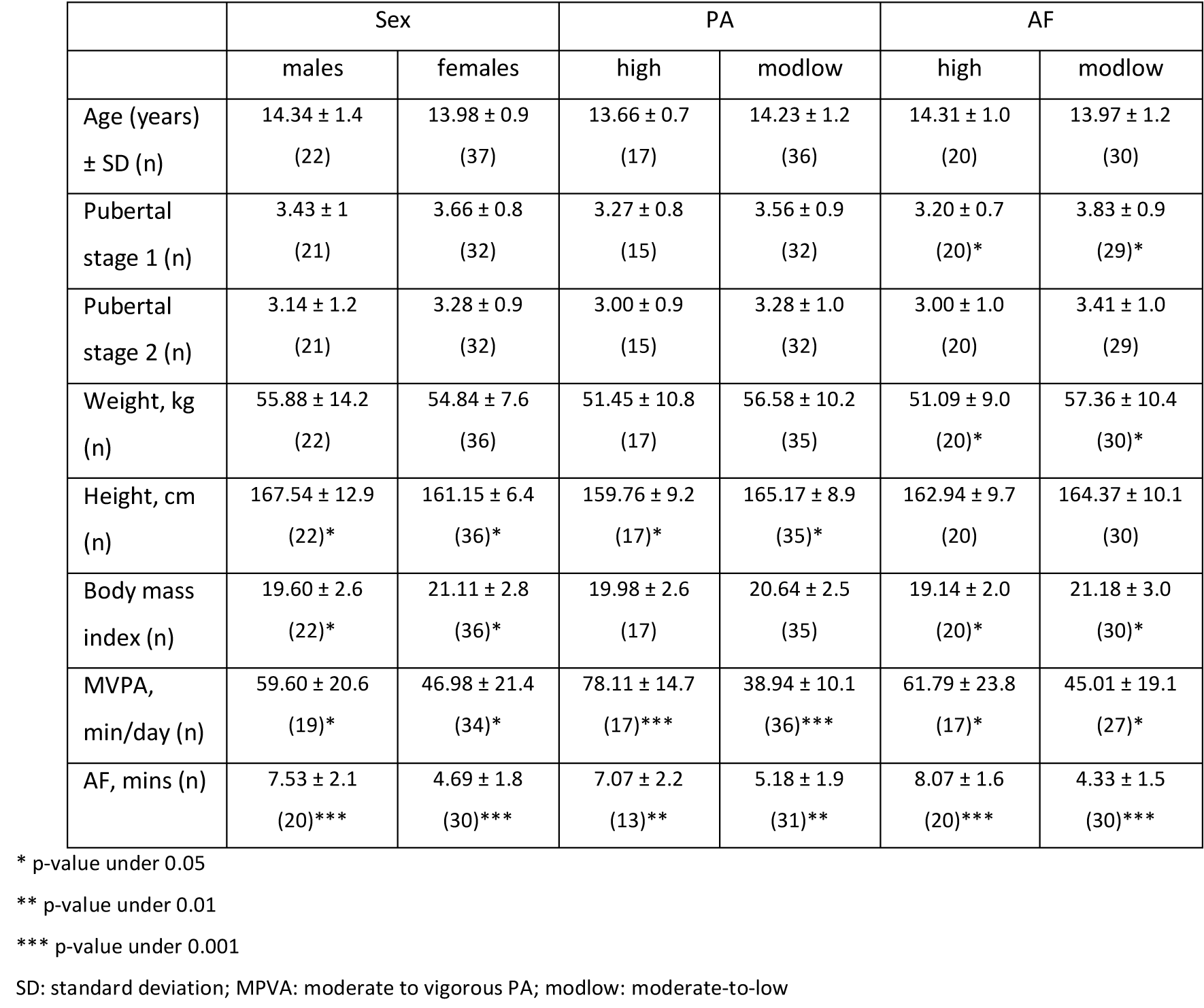
Demographic information and statistical differences for the groups based on physical activity (PA), aerobic fitness (AF) and sex.

|  | Sex |  | PA |  | AF |  |
| --- | --- | --- | --- | --- | --- | --- |
|  | males | females | high | modlow | high | modlow |
| Age (years)<br>± SD (n) | 14.34 ± 1.4<br>(22) | 13.98 ± 0.9<br>(37) | 13.66 ± 0.7<br>(17) | 14.23 ± 1.2<br>(36) | 14.31 ± 1.0<br>(20) | 13.97 ± 1.2<br>(30) |
| Pubertal<br>stage 1 (n) | 3.43 ± 1<br>(21) | 3.66 ± 0.8<br>(32) | 3.27 ± 0.8<br>(15) | 3.56 ± 0.9<br>(32) | 3.20 ± 0.7<br>(20)* | 3.83 ± 0.9<br>(29)* |
| Pubertal<br>stage 2 (n) | 3.14 ± 1.2<br>(21) | 3.28 ± 0.9<br>(32) | 3.00 ± 0.9<br>(15) | 3.28 ± 1.0<br>(32) | 3.00 ± 1.0<br>(20) | 3.41 ± 1.0<br>(29) |
| Weight, kg<br>(n) | 55.88 ± 14.2<br>(22) | 54.84 ± 7.6<br>(36) | 51.45 ± 10.8<br>(17) | 56.58 ± 10.2<br>(35) | 51.09 ± 9.0<br>(20)* | 57.36 ± 10.4<br>(30)* |
| Height, cm<br>(n) | 167.54 ± 12.9<br>(22)* | 161.15 ± 6.4<br>(36)* | 159.76 ± 9.2<br>(17)* | 165.17 ± 8.9<br>(35)* | 162.94 ± 9.7<br>(20) | 164.37 ± 10.1<br>(30) |
| Body mass<br>index (n) | 19.60 ± 2.6<br>(22)* | 21.11 ± 2.8<br>(36)* | 19.98 ± 2.6<br>(17) | 20.64 ± 2.5<br>(35) | 19.14 ± 2.0<br>(20)* | 21.18 ± 3.0<br>(30)* |
| MVPA,<br>min/day (n) | 59.60 ± 20.6<br>(19)* | 46.98 ± 21.4<br>(34)* | 78.11 ± 14.7<br>(17)*** | 38.94 ± 10.1<br>(36)*** | 61.79 ± 23.8<br>(17)* | 45.01 ± 19.1<br>(27)* |
| AF, mins (n) | 7.53 ± 2.1<br>(20)*** | 4.69 ± 1.8<br>(30)*** | 7.07 ± 2.2<br>(13)** | 5.18 ± 1.9<br>(31)** | 8.07 ± 1.6<br>(20)*** | 4.33 ± 1.5<br>(30)*** |

\* p-value under 0.05
\*\* p-value under 0.01
\*\*\* p-value under 0.001
SD: standard deviation; MPVA: moderate to vigorous PA; modlow: moderate-to-low

Males showed higher levels of MVPA (moderate to vigorous PA, as measured by the accelerometer on a daily basis) than females (males: 59.60 ± 20.66 min/day; females: 46.98 ± 21.4 6 min/day) (t[51] = 2.083, p = 0.042) (see Table 1). Males also showed higher levels of AF (minutes until exhaustion in maximal shuttle run test) (males: 7.53 ± 2.12 min; females: 4.69 ± 1.84 min) (t[48] = 5.024, p < 0.001) (see Table 1).

PA and AF showed a clear association (r = 0.524, p < 0.001) in the whole sample (see Fig. 1A and Experimental Procedures in section 3.2 for details about the subject groups). When females and males were analyzed separately, this association was significant for males (r = 0.577, p = 0.015) but not for females.

**Fig 1.**
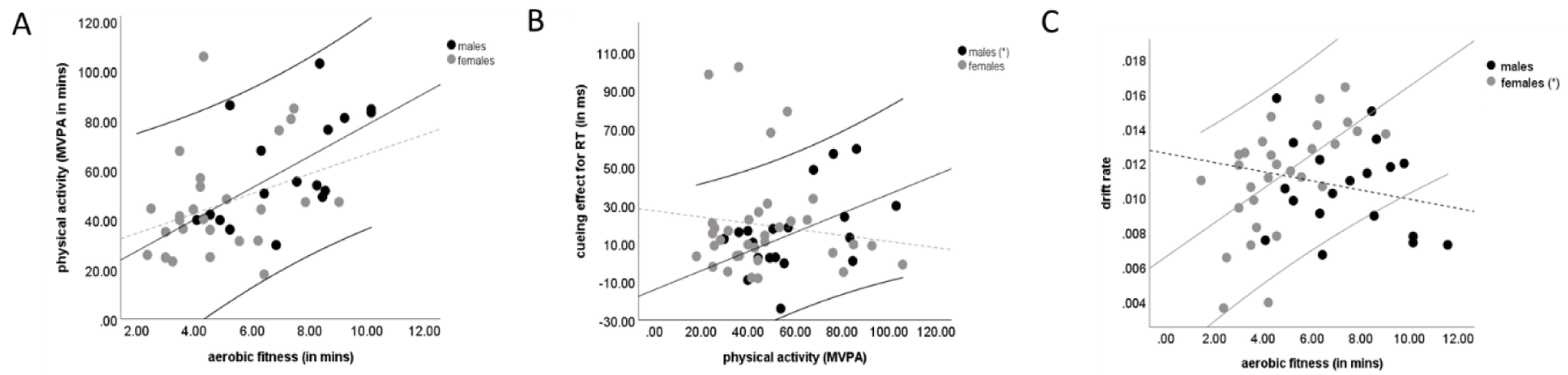
Associations of studied variables. Black dots represent males and grey dots represent females. The solid central line represents the linear trend line and the two external lines represent the 95% confidence interval for the significant sex group. The dotted central line represents the linear trend line for the non-significant sex group. (A) Aerobic fitness (maximum time during maximal shuttle run test, min) plotted against physical activity (MVPA, moderate to vigorous physical activity, min/day) for males and females. Solid trend line and confidence interval for males, dotted trend line for females. (B) Cueing effect for reaction times from visuospatial covert attention task plotted against physical activity for males and females. Solid trend line and confidence interval for males, dotted trend line for females. (C) Drift rate from visuospatial covert attention task plotted against aerobic fitness for males and females. Solid trend line and confidence interval for females, dotted trend line for males.

### Associations between physical activity, aerobic fitness and attention/inhibition skills

During the visuospatial covert attention task, a cue (a small fish, valid in 75% of the cases) was presented before two possible targets (two sharks on each side of the screen). Participants were instructed to answer as fast as possible which shark (located in the left or right hemifield) opened its mouth more to eat the small fish (see Fig. 2A). The task requirements, and the underlying oscillatory dynamics in the left and right hemispheres (based on extensive literature [Jensen and Mazaheri, 2010; Bonnefond and Jensen, 2013; Vollebregt et al., 2015]), are schematically illustrated in Fig 2B. From this task, RT, cueing effect for RT, cueing effect for accuracy and EZ model variables (drift rate, boundary separation and non-decision time) were included in statistical analysis. Besides this covert attention task (MEG-task), attention and inhibitory control were measured using a behavioral test battery including a modified Eriksen Flanker task and a reaction time (RTI) test.

**Fig 2.**
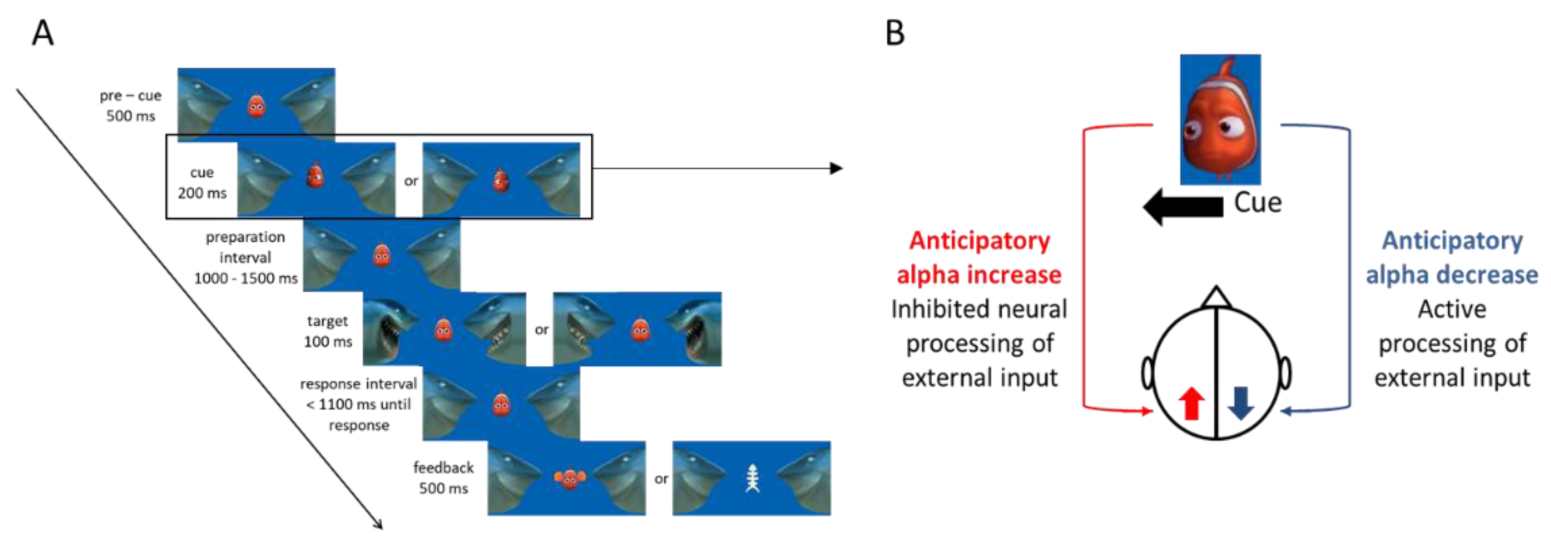
Schematic representation of Posner’s modified visuospatial covert attention task. (A) The progress of the task: a visual cue (fish looking either left or right) was followed by a target (one shark opens its mouth more than the other). Participants should detect the target as soon as possible by pressing the left or right button. (B) Illustration of the alpha modulation hypothesis (Jensen and Mazaheri, 2010) and the expected change in alpha power during the visuospatial covert attention task in the two hemispheres.

### Physical activity vs. performance in visuospatial covert attention task (MEG task)

Visuospatial covert attention task measures did not correlate with PA levels. When this correlation was examined separately for males and females, the cueing effect for RT (reaction time for uncued minus reaction time for cued targets) correlated with PA levels (MVPA, see Experimental Procedures, section 3.2) in males (r = 0.485, p = 0.035), showing that the higher the PA level, the higher the difference between RTs for uncued minus cued targets (see Fig. 1B). No significant correlations were observed for females.

### Aerobic fitness vs. performance in visuospatial covert attention task (MEG task)

Visuospatial covert attention task measures did not correlate with AF levels. In the separate analysis for males and females, significant correlations were found between drift rate and levels of AF for females (r = 0.580, p = 0.001), showing that higher AF was associated with a higher drift rate (see Fig. 1C). No significant correlations between EZ-model variables and AF were found for males.

### Physical activity and aerobic fitness vs. performance in attention and inhibitory control tests (behavioral tests outside MEG)

Flanker variables (RT for compatible conditions) and RTI measures (RT for five-choice condition, movement time for five-choice condition) included in the analysis were not significantly correlated with PA or AF measures.

Flanker variables (RT for compatible conditions; the number of errors in compatible incongruent conditions) and RTI measures (RT for five-choice condition, movement time for five-choice condition) included in the analysis were not significantly correlated with visuospatial covert attention task measures.

### Associations between physical activity, aerobic fitness and brain measures

The temporal variation of the spectral power at the alpha band in each hemisphere was measured in cued (i.e. cue to the left hemifield for the left hemisphere target and cue to the right hemifield in the right hemisphere target) and uncued (i.e. cue to the left hemifield for the right hemisphere target and cue to the right hemifield for the left hemisphere target) conditions. To examine the modulation of alpha oscillations in response to the cue, left cued vs. right uncued and left uncued vs. right cued conditions were contrasted. Modulation indexes (MI) for each hemisphere were calculated as the power from left cued trials minus right cued trials normalized by their mean (Vollebregt et al., 2015).

Average time-frequency representations (TFRs) over selected occipital MEG sensors in each hemisphere (see Fig. 3B) were obtained. MI for left and right hemispheres were used to test the existence of alpha modulation (i.e. increase vs. decrease in response to the cue) and differences between groups. Three different time-windows were selected *a priori* and used for further analysis (0–400 milliseconds [ms], 401–800 ms and 801–1,200 ms post-cue). Repeated measures ANOVA (performed separately for PA and AF groups) were completed in order to test the differences in brain measures (left increase and right decrease of alpha MI in the three time-windows).

**Fig 3.**
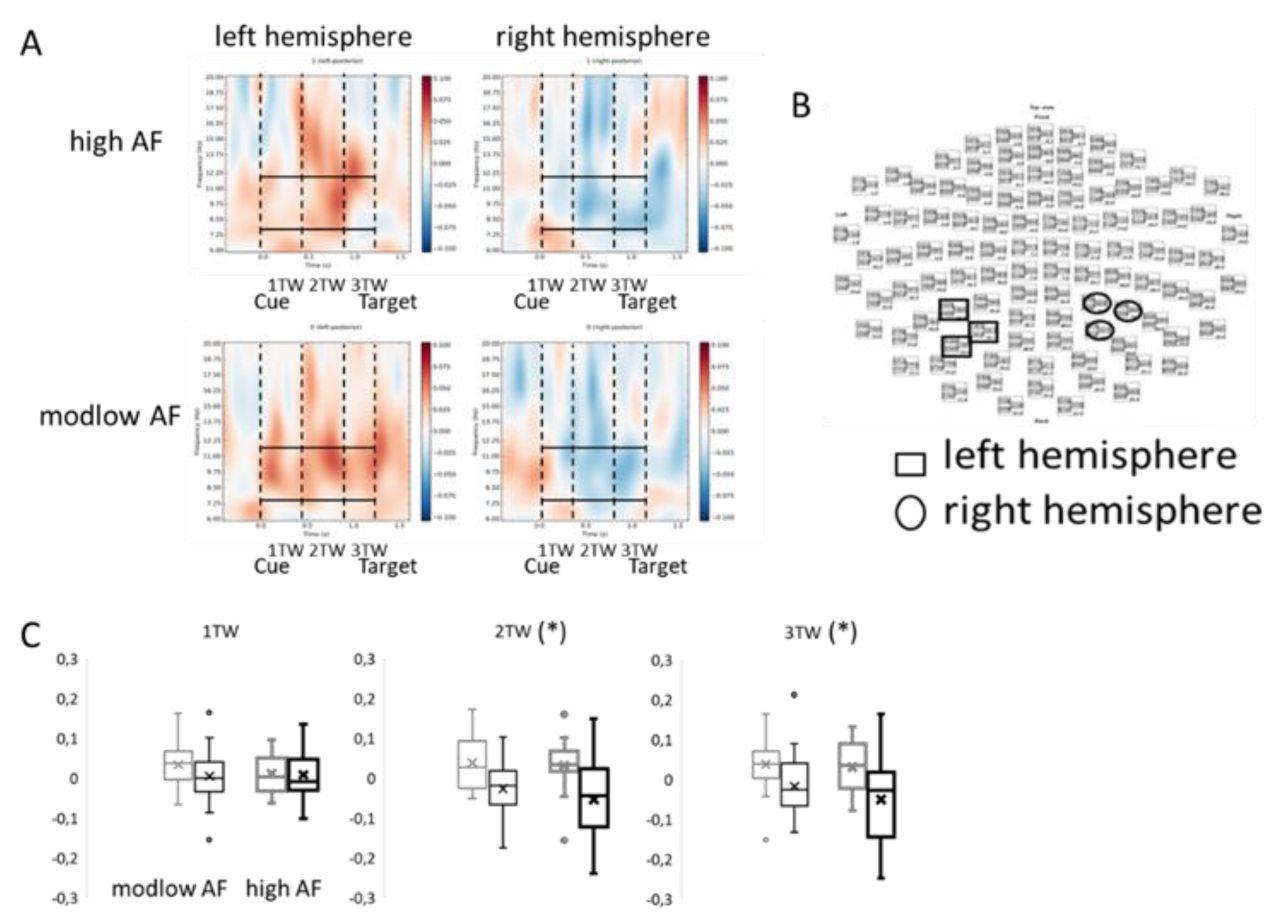
Modulation of alpha power in response to the spatial cue for AF groups. (A) Time-frequency representations of the alpha power modulation index in the left and right hemispheres for moderate to low (modlow) and high aerobic fitness (AF) groups. Dotted lines delimit the three *a priori* selected time-windows. The first dotted line corresponds with the onset of the cue and the last dotted line corresponds with the onset of the target. Solid horizontal lines delimit the alpha rhythm frequency range (8–12 Hz). (B) Location of selected channels for calculating the alpha modulation index in the left and right hemispheres (square for the left hemisphere and circle for the right hemisphere. (C) Boxplot representation of left (in grey) and right (in black) hemispheres in each time-windows for modlow (light color) and high (dark color) AF groups.

The presence of post-cue alpha modulation was verified. As the MI was calculated by subtracting the right cued from the left cued conditions (divided by their sum), and the alpha increases ipsilaterally to the attended hemifield, MI was expected to be positive in the left hemisphere and negative in the right hemisphere (see Fig. 2B). Figure 3A illustrates the cue-induced modulation of spectral power in left and right hemispheres (time-frequency representation) for AF groups. In the same way, Figure 4A show the time-frequency representation obtained for PA groups. Both rANOVAs showed a clear main effect of hemisphere (AF as between subjects factor: hemisphere F_(1,48)_ = 19.114 p < 0.001; PA as between subjects factor: hemisphere F_(1,51)_ = 17.246 p < 0.001) in alpha power reflecting the expected pattern between hemispheres (left > right).

**Fig 4.**
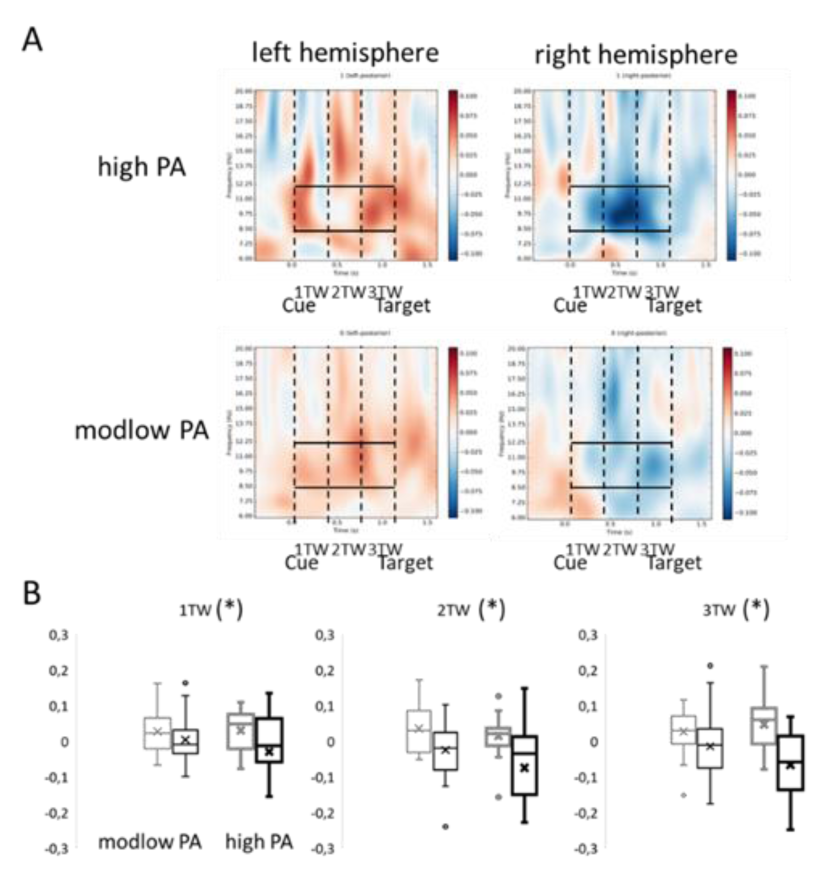
Modulation of alpha power in response to the spatial cue. (A) Time-frequency representations of the alpha power modulation index in the left and right hemispheres for moderate to low (modlow) and high physical activity (PA) groups. Dotted lines delimit the three *a priori* selected time-windows. The first dotted line corresponds with the onset of the cue and the last dotted line corresponds with the onset of the target. Solid horizontal lines delimit the alpha rhythm frequency range (8–12 Hz). (B) Boxplot representation of left (in grey) and right (in black) hemispheres in each time-windows for modlow (light color) and high (dark color) PA groups.

When AF was used as between subjects factor no significant interaction or between subjects effect were found for AF. A significant interaction of hemisphere x time-windows was found (hemisphere x TW F_(2,96)_ = 8.339 p < 0.001). A main effect of time-window was close to significance (TW: F_(1.543, 74.071)_ = 3.306 p = .054). A follow-up test (separately for each time-window), conducted in order to clarify the more detailed differences, revealed a clear main effect of hemisphere in alpha power in the second (hemisphere: F_(1,48)_ = 20.962 p < 0.001) and third (hemisphere: F_(1,48)_ = 17.104 p < 0.001) time – windows reflecting the left > right alpha modulation. The direction of the differences between hemispheres can be seen in Figure 3C.

When PA was used as a between subjects factor, there was a tendency towards significant main effect for group (TW: F_(1, 51)_ = 3.544 p = .065). Moreover, also the effect of time-window (TW: F_(1.727, 88.099)_ = 2.983 p = .063) and hemisphere x time-window interaction (hemisphere x TW: F_(1.728, 88.152)_ = 2.474 p = .098) showed small p-values, however not reaching the 0.05 level of significance. To perform similar analysis as for AF groups, and to test our hypotheses on group differences, we further tested the effect of hemisphere and group in each time-window. The follow up test revealed a main effect of hemisphere in the three TWs (1TW: F_(1, 51)_ = 4.831 p = .033; 2TW: F_(1, 51)_ = 14.838 p < 0.001; 3TW: F_(1, 51)_ = 19.580 p < 0.001) and a hemisphere x PA group interaction for the third TW (hemisphere x TW: F_(1, 51)_ = 4.268 p = .044). Also, a main effect of PA group in the second time-window (F (_1,51_) = 4.468 p = .039) was found, probably reflecting the way alpha MI is calculated. The direction of the differences between hemispheres can be seen in Figure 4B.

### Associations between MEG measures and cognition

In order to test the correlation between the brain activation and behavioral performance, interhemispheric asymmetry (the difference between left – right hemisphere alpha MIs) was calculated as a single variable indexing the observed difference between hemispheres.

The interhemispheric asymmetry during the third time-window showed a significant correlation with drift rate. A bigger interhemispheric asymmetry during the third time-window was associated with lower drift rate values (r = - 0.359, p = 0.006) (see Fig. 5B). No significant correlations were identified between the interhemispheric asymmetry during the third time-window and other behavioral measures from the MEG task, the attention test (RTI) or the inhibitory control task (Flanker).

**Fig 5.**
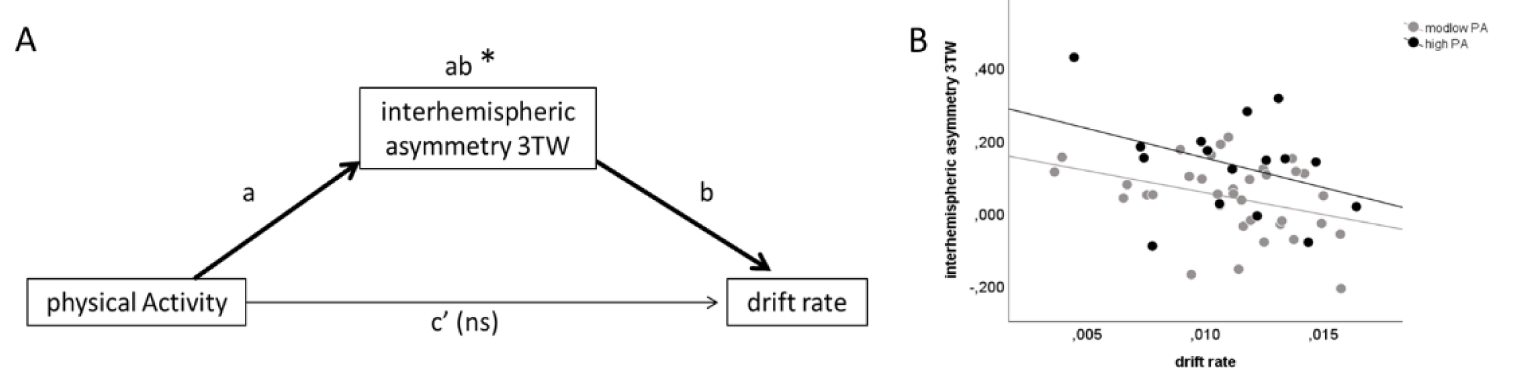
Schematic illustration of the mediation model. (A) Mediation path examined about the role of interhemispheric asymmetry of alpha MI during the third time-window in mediating the relationship between physical activity and drift rate from the visuospatial covert attention task. Darker arrows indicate a significant path (only an indirect effect). (B) Interhemispheric asymmetry during the third time-window plotted against drift rate for both physical activity groups. Black dots and trend line correspond to high physical activity group and grey color dots and trend line correspond to moderate-to-low (modlow) physical activity group.

### Brain oscillatory dynamics as mediators between physical activity or aerobic fitness and cognition

The possible mediator role of brain oscillatory measures in the relationship between PA or AF and cognition was tested. A mediator analysis was used to test the underlying assumption that task-related modulation of oscillatory power mediates the influence of PA or AF on performance in visuospatial covert attention task. First, significant correlations between brain measures, PA or AF groups and cognitive variables were tested. Of the brain measures, only the interhemispheric asymmetry of alpha power during the third time-window fulfilled this assumption correlating with PA groups (but not AF groups) and cognitive variables (only drift rate). The resultant combination was tested with a mediator model using MPlus software by means of a bootstrap of 1,000 samples. Mediation was tested by determining whether the confidence interval for the indirect effect contained zero (Fritz and MacKinnon, 2007). The intervals not including zero were considered significant.

The mediation model was built with PA groups as the independent variable, drift rate as the dependent variable and interhemispheric asymmetry during the third time-window as the mediator. Indirect effect was significant (ab estimate = – 0.107, bias-corrected bootstrap confidence interval = [– 0.306; 0.000]) but direct and total effects were not (direct: c’ estimate = 0.104, p = 0.458; total: c estimate = – 0.003, p = 0.983) (see Fig. 5A). In this model, the relationship between independent (PA groups) and dependent (drift rate) variables was not significant, but this relationship was increased in magnitude when late interhemispheric asymmetry was included as a mediator. As the mediated effect and the direct effect have opposite signs, the model indicates substantial mediation (MacKinnon, 2008). The results of this model thus suggest that the late interhemispheric asymmetry of alpha mediates the association between PA levels and drift rate in the visuospatial covert attention task across the whole sample.

To determine if sex influenced the above mediation effect, a moderation in the mediator effect analysis was performed. The results showed no significant difference in indirect effects between males and females (d4 estimate = – 0.103, p = 0.798, bias-corrected bootstrap confidence interval = [– 1.144; 0.503]). The results of this model suggest that sex does not moderate the indirect or total effects of late interhemispheric asymmetry of alpha in the association between PA levels and drift rate.

## DISCUSSION

This study evaluated the influence of physical activity and aerobic fitness on brain oscillatory measures underlying anticipatory attention allocation within the same sample of adolescents. As expected, even though the levels of physical activity and aerobic fitness were strongly correlated, they showed specific associations with cognitive performance and underlying brain measures. Adolescents, especially males, with higher levels of physical activity showed stronger utilization of the cue in anticipating their response, reflected as bigger cueing effect (RT difference between uncued vs. cued targets) in reaction time. In line with these behavioral findings, the higher physical activity group showed stronger cue-induced interhemispheric alpha asymmetry during the allocation of attention in a modified Posner’s cueing task, which engages selective visuospatial attention vs. inhibition for the opposite hemifields. The exploratory mediation analysis suggested that this interhemispheric alpha asymmetry mediated the association between physical activity status and drift rate, a measure of information processing speed, calculated as an individual performance measure in the Posner’s cueing task. Larger cue-induced asymmetry was associated with slower overall information processing at the behavioral level, in an analysis integrating valid and invalid spatial cues. Although males and females showed a differential association between physical activity and aerobic fitness and cognition, this mediator effect was not influenced by sex.

Males with a higher level of physical activity indicated stronger cue-based anticipation, as the bigger difference in reaction times between cued and uncued trials, suggesting a greater reliance on the cue. This result is partly supported by an earlier study showing a complex association between attentional control and physical activity measures especially in males (Booth et al., 2013). By measuring selective attention, sustained attention and attentional control/switch, Booth and colleagues found that total volume of PA (mostly including light intensity activity) predicted lower performance on the attention tasks, while higher MVPA was associated with better executive function performance in adolescent males. In contrast, Vanhelst et al. (2016) reported a positive effect of physical activity on attention capacity in adolescents independent of sex. The possible sex difference indicated by our findings needs to be confirmed by further studies, as we had fairly small number of participants (especially when both sex and physical activity groups were separately analyzed). Differences between our results and earlier findings might be related to the way the attentional tests were administered (computerized vs. paper and pencil), and to the way cueing was used in the experiments. In our study, we used rather high ratio of cued/uncued trials (75/25 %) which may emphasize attentional shifting over efficient attention control for optimal performance. In sum, there is some evidence for an association between higher levels of physical activity and more dependence on the cue in adolescent males than in females. Interestingly, we demonstrated a modest but significant difference at the brain level, related to the cue-based allocation of attentional resources. The significant hemisphere x physical activity group interaction indicated a larger difference between the ipsilateral increase and contralateral decrease of alpha oscillations in the two hemispheres. This effect appeared at around 800 ms after the cue onset, right before the target. This means that for adolescents with higher level of physical activity, the cue-based engagement of visual attentional resources (inhibition of the uncued and attending to the cued hemifields) is stronger than for individuals with lower levels of physical activity. This result can be interpreted according to Banich (1998), who suggested that, depending on task demands, the interaction between the brain hemispheres, rather than the specific processes accomplished by each of them, can influence selective attentional functioning. Therefore, physical activity seems to be related to relying more on cues for engaging anticipatory attentional control, reflected in both behavioral measures (reaction time difference between uncued vs. cued targets) and brain measures (anticipatory interhemispheric asymmetry of alpha oscillations). Indeed, earlier studies have indicated individual differences in the adherence to the cue and its subsequent effect in target processing and the ultimate behavioral performance (Hong et al., 2020).

The effect of physical activity was not observed in the overall performance in the selective attention task, it was seen instead in the strategy of performing the task, perhaps reflecting prioritization of anticipatory focusing of attentional resources over adaptive shifting of attention. Indeed, the overall behavioral outcome did not differ, but only the reliance on the cue. In fact, as evidenced by the subsequent mediation analysis, the strong interhemispheric asymmetry was linked with less efficient attentional processing, as measured by the drift rate variable (reflecting speed of information processing), when a single measure of drift rate was calculated, integrating cued and uncued targets. Our results would thus suggest that rather than focusing on the absolute benefit of physical activity on cognitive functions, it might be useful to examine the influence of physical activity on the more specific mechanisms of information processing in the brain. Indeed, it might be that by specifically influencing part of the neurocognitive resources underlying attentional control, physical activity may appear beneficial for executive functions in some tasks but not in others. Thus, differential effects across tasks could depend on the specific neurocognitive requirements of a given task. This interpretation of our data would help to explain the discrepant findings concerning the effects of physical activity on executive functions in youth (Alvarez-Bueno et al., 2017; Esteban-Cornejo et al., 2015).

What could then be the underlying mechanisms by which physical activity contributes to the neural bases of attention? Our experimental manipulation focused on the attentional control that requires communication between the left and right visual areas. The dynamic interaction between hemispheres, modulating the processing capacity of the brain, is proposed to occur via the corpus callosum (Banich, 1998; Qin et al., 2016). A recent study by our group (Ruotsalainen et al., 2020) investigated the connection between both aerobic fitness and physical activity with the white matter in adolescents using MRI and fractional anisotropy (assessing microstructural changes in the brain’s white matter). Although the main results in this study showed that only the level of aerobic fitness was related to white matter properties, it was also described that the level of fractional anisotropy in the body of the corpus callosum could moderate the relationship between physical activity and working memory. High levels of physical activity were positively related to working memory only when the fractional anisotropy level was low (which could occur in the earlier stages of corpus callosum maturation). The authors suggest that specific tracts of white matter (like in the case of corpus callosum) could moderate the relationship between both physical activity and aerobic fitness with cognition in adolescents. If the changes in white matter fractional anisotropy is associated with variations in alpha oscillatory activity, as it is suggested by some studies (Jann et al., 2012; Valdés-Hernández et al., 2010), it would be critical to acknowledge the maturational state (and consequently, the level of plasticity) of the brain to explain the possible effects of physical activity and fitness over cognitive performance. Further research is warranted to clarify the role of interhemispheric interaction via the corpus callosum in the influence of physical activity and aerobic fitness over executive functions (such as attentional control/inhibition and working memory).

Our exploratory mediation analysis indicated that the interaction (or, more specifically, the degree of imbalance) between the hemispheres (interhemispheric asymmetry) during attention allocation seemed to mediate the relationship between physical activity levels and performance in a selective attention task. Physical activity increased the imbalance between the ipsi- and contralateral hemispheres according to the selective attention task requirements. However, this increment in interhemispheric asymmetry did not improve the efficiency in the selective attention task (as measured by drift rate). This unexpected result could be partly explained by a strategy of choice emphasizing the processing of the cue. Adolescents with higher physical activity could be prioritizing the anticipation for upcoming information, as was hinted also by the stronger cueing effect. Similar results were obtained in a longitudinal study measuring the association of preadolescent children’s motor competences with working memory maintenance and neurophysiological measures of task preparation during 9 months (Ludyga et al., 2020). Ludyga et al., (2020) described a more effective utilization of the cue-relevant information by an increase in the cue-P300 amplitude during the task preparation stage in a Stenberg paradigm (used to assess working memory capacity) in children with high motor competences, thus supporting our findings. The authors suggest that children with high motor competence exhibited a more proactive control strategy allowing active maintenance of task goals. It seems that these differences in strategy between children with low and high accelerometer-measured motor competences can also be observed in adolescents when solving a selective attention task. However, this strategy could have a cognitive cost leading to a worse global efficiency in the selective attention task, especially when the proportion of uncued targets in the task is rather high. The cost-benefit effect of relying on the cue could also arise due to shared neural resources between selective attention and working memory (For a review see Gazzaley and Nobre, 2012). Physical activity has been associated with increased working memory capacity in children and adults (Alvarez-Bueno et al., 2017; Kamijo et al., 2011; Ludyga et al., 2020). In a task requiring both, working memory (holding the side of the target in memory) and possible attentional switching, the strategy relying on working memory may limit the resources for adaptive attentional switching.

In line with earlier studies in youth, aerobic fitness and physical activity were found to be strongly correlated (Butte et al., 2007; Gutin et al., 2005; Ruotsalainen et al., 2019). Nevertheless, they showed different associations with executive control and its underlying brain processes. Although the aerobic fitness has been shown to be more strongly associated with brain structural measures than physical activity (Ruotsalainen et al., 2019; Ruotsalainen et al., 2020), we did not find any relationship between aerobic fitness and brain oscillatory dynamics underlying anticipatory attention allocation. However, in behavioral measures higher aerobic fitness was related to higher drift rate values in the selective attention/inhibition task. Interestingly, this relationship was observed only in females. Drift rate was calculated based on the variance of reaction times for correct decisions and the proportion of correct decisions as part of the EZ diffusion model for two-choice RT tasks (Wagenmakers et al., 2007). Thus, the drift rate has been interpreted as an index of processing speed in decision tasks (Wagenmakers et al., 2007). Previous studies have also reported the association between fitness and attention/inhibition in adolescents (Cadenas-Sanchez et al., 2017; Huang et al., 2015; Marchetti et al, 2015; Westfall et al., 2018). Thus, aerobic fitness seems to be related with the selective attention/inhibition processing but probably the anticipatory gating of relevant information, reflected as asymmetric alpha oscillations in visual areas is not specifically influenced by aerobic fitness.

The selective advantage of aerobic fitness for females’ cognition/executive functions has been reported before for older adult populations (Baker et al., 2010; Barha et al., 2017; Colcombe and Kramer, 2003; van Uffelen et al., 2008). Our findings support and expands previous studies by showing a sex-dependent positive effect of aerobic fitness over selective attention/inhibition also earlier in life. There are some potential underlying explanations for the observed female-specific advantage of fitness for executive functions. Based on a study exploring the association between circulating BDNF levels and cognitive functioning, it has been hypothesized that the brain-derived neurotrophic factor (BDNF) is more closely related to cognition in women compared to men (Komulainen et al., 2008). Moreover, some studies have suggested that increased feminine hormone, estradiol, might be related to greater BDNF expression in the cortex and hippocampus of females of different species (Scharfman et al., 2003; Singh et al., 1995; Sohrabji et al., 1995). More studies are needed to investigate the possible mediator role of sex hormones in the relationship between aerobic fitness-related increments of BDNF and cognition.

Although our findings need to be replicated with bigger sample sizes, our results suggest that the associations between physical activity, aerobic fitness and (neuro)cognitive measures are partly dependent on sex. The aerobic fitness level in females seemed to be associated with more efficient attentional/inhibitory processing (drift rate) in the selective attention/inhibition task. On the other hand, physical activity level in males seemed to be associated with the clearer benefit of cued over uncued targets (presumably reflecting a choice of strategy). It is important to note that we cannot rule out the effect of other sex-related variables, such as peer group influence, personality and preferences in free-time activities, that are likely to influence physical activity more than influencing aerobic fitness (Palmer-Keenan and Bair, 2019; Sevil et al., 2018), which thus impacts the interpretation of our results. In line with earlier studies (Booth et al., 2013; Tomkinson et al., 2018; Woll et. al., 2011), physical activity levels were higher in males than in females. It may be that higher levels of activity in boys at this age also reflect engagement to regular, goal-directed training, which could be generalized as higher competitiveness, also in other contexts. This could influence the choice of strategy in the visuospatial attention task. In the same way, underlying genetic factors could influence the association between aerobic fitness and attentional processing in females.

Our results, showing a mediator effect of brain dynamics on the association between physical activity and selective attention, also raise the probable developmental perspective in the study of the underlying brain mechanisms in the association between physical activity or aerobic fitness and attention/inhibition. Benefits from physical activity have been reported mostly using behavioral and neuroelectric techniques (ERPs) for young adults (Hillman et al., 2006; Kamijo et al., 2011; Kamijo and Takeda, 2009). However, aerobic fitness seems to benefit selective attention and inhibition, especially during childhood (Chaddock et al., 2011; Pontifex et al., 2011; Voss et al., 2011) and older adulthood (Colcombe et al., 2004; Prakash et al., 2011). Evidence of benefits from physical activity on attention/inhibition is weaker in other age groups. Altogether, this suggests a U-shaped relationship between physical activity and selective attention/inhibition across the lifespan. Further work is required to establish whether this really is the case or whether this reflects only an insufficient number of studies tackling regular physical activity engagement across the lifespan.

A major limitation of this study resides in its small sample size, which limits the strength of our interpretations. Another limitation lies in the cross-sectional nature of this study, which prevents us from making causal inferences from the associations found for physical activity or aerobic fitness. The results of our mediator analysis provide valuable insights for understanding brain processes that might mediate the effects of physical activity on cognition, although they should be interpreted with caution. The absence of direct effect in our mediation model (direct influence of physical activity over drift rate) could be a result of the reduced sample size. It may thus be that there was sufficient power to detect the indirect effect, but insufficient power to detect the direct and total effects. Longitudinal studies would be needed for a better understanding of the neural level mediators of the cognitive benefits of physical activity and aerobic fitness during different phases of life. Finally, this study measured only physical activity and aerobic fitness effects over selective attention and inhibitory control processes in visual modality. Testing these effects in different modalities or cross-modal interactions should also be considered in future research. We failed to replicate the associations between aerobic fitness and reaction times/accuracy in the Flanker task (Westfall et al., 2018). Our participants showed a ceiling effect in accuracy for this task, possibly influencing the reliability of this task measure in our experiment. On the other hand, the absence of a link between aerobic fitness and inhibitory control as measured by the Flanker task in adolescents was also reported by Stroth et al. (2009), thus supporting our findings.

In conclusion, our results showed that higher levels of physical activity, but not aerobic fitness, were related to improved anticipatory brain processes underlying allocation of attention in the adolescent brain. However, the anticipatory interhemispheric asymmetry was related with reduced overall performance in the selective attention task, thus suggesting a choice of strategy prioritizing the cue processing. Female and male adolescents showed dissociable effects of physical activity and aerobic fitness on anticipatory selective attention/inhibitory control.

## EXPERIMENTAL PROCEDURES

### Participants

Sixty-three adolescents were recruited to participate from among the participants of a larger study related to the Finnish Schools on the Move Program (Joensuu et al., 2018; Syväoja et al., 2019) measuring the behavioral effects of PA and AF on cognition. The potential subjects were informed about the study and voluntarily manifested their interest to participate. All subjects were native Finnish speakers and came from the Central Finland area (n = 54) or the South Finland area (n = 9). Sixty participants were classified as right-handed and three as left-handed by the Edinburgh Handedness Inventory. From the initial 63 participants, four were excluded from further analysis due to low quality of MEG data (n = 2) or very few responses in the uncued condition during the visuospatial covert attention task (n = 2). Finally, the analysis was done with a total of 59 participants (22 males and 37 females) with age ranges from 12.8 to 17.0 years old (14.11 ± 1.07 years).

All participants were volunteers, and they as well as their legal guardians signed informed consent before the beginning of the study in agreement with prior approval of the Central Finland Healthcare District Ethical Committee. Participation in this study was compensated with a €30 gift card. The sample did not include participants with neurological disorders, major medical conditions or those using medication that influences the central nervous system. All participants had normal or corrected to normal vision.

Self-reports about the participants’ stages of puberty with the Tanner Scale (Marshal and Tanner, 1969; 1970) were used to measure the pubertal development in our sample. All the participants reported being between categories 2 and 5 for pubertal stage 1 and between 1 and 5 for pubertal stage 2. For demographic information and statistical differences for the groups based on AF, PA or sex, see Table 1.

### Measures of physical activity and aerobic fitness

AF was measured with the shuttle run test (Leger et al., 1988; Joensuu et al., 2018), a measure widely used to estimate a person’s maximum oxygen uptake (VO_2_max). Participants were instructed to run between two lines with a separation of 20 m. Speed should have accelerated every time they heard an audio signal. The time participants spent before failing to reach the end lines in two consecutive tones indicated their level of aerobic fitness. The speed in the first and second levels was 8.0 and 9.0 km/h, respectively. Afterward, speed increased by 0.5 km/h per level. The duration of each level was one minute. The number of minutes that each participant lasted until exhaustion and abandoning the test (normalized by gender and age) was used to measure the participant’s AF level. To differentiate them by their level of AF into high or modlow AF groups, the distribution of minutes until exhaustion was divided into three equal parts. Participants in the highest tertile (cutoff point tagging 66% of the distribution) were considered as being part of the high AF group. In the same way, participants in the two lowest tertiles were allocated to the modlow AF group.

PA was measured with accelerometers, specifically triaxial ActiGraph (Pensacola, FL, USA) GT3X+ and wGT3X+ monitors, which the participants wore during seven consecutive days (Joensuu et al., 2018). They were instructed to wear it on their right hip during waking hours except for water-related activities. A valid measurement day consisted of at least 10 h of data. Data were collected in raw 30 Hz acceleration, standardly filtered and converted into 15 s epoch counts. Periods of at least 30 min of consecutive zero counts were considered as non-wear time. A customized Visual Basic macro for Excel software was used for data reduction. A cut-off point of ≥2296 cpm (Evenson et al., 2008) was used to define moderate to vigorous intensity PA (MVPA, min/day). MVPA was calculated as a weighted mean value of MVPA per day ([average MVPA min/day of weekdays × 5 + average MVPA min/day of weekend × 2] /7). To differentiate participants by their PA level into the high or modlow PA groups, the distribution of MVPA (min/day) was divided into three equal parts. Participants in the highest tertile (cutoff point tagging 66% of the distribution) were placed into the high PA group. In the same way, participants in the two lowest tertiles were allocated to the modlow PA group.

PA was measured in a total of 53 participants and AF was measured in a total of 50 participants. School absence on the day of the test was the reason for missing AF values (2 males and 7 females), while an insufficient number of valid measurement days (two weekdays and one weekend day) was the reason for missing data of regular PA measurements (3 males and 3 females). All subjects had at least one measure of PA or AF.

### Stimuli and task

We used a visuospatial covert attention paradigm from Vollebregt et al. (2015) based on a modified Posner’s cueing paradigm (Posner, 1980). The task of the participants was to save a fish from being eaten by a shark. A schematic representation of the sequence of phases displayed in the task is shown in Fig. 2A. The first phase was the pre-cue period (500 ms), when the subjects were presented with a fish in the middle of the screen as a fixation point and two sharks on both sides of the screen. Next, a cue was presented during 200 ms consisting of the fish’s eyes looking to the left or the right shark. The cue indicated the side of the screen where the target would appear. The target (both sharks opened their mouths but one more than the other) was presented after 100 ms. The probability of validly cued targets (the gaze of the fish was directed towards the same side as the targeted shark) was 75%. Afterward, participants had a preparation interval varying in time (1,000–1,500 ms) to avoid the subject’s prediction of the target’s occurrence. The duration of the response interval following the target presentation was determined by the subject’s response with a maximum time of 1,100 ms. Subjects had to report (by using their index finger in a MEG-compatible response pad) the side corresponding to the shark that opened its mouth more. Afterward, the feedback (a happy fish for correct responses or a fishbone for errors) was presented. During the whole task, subjects were instructed to continuously watch the fish’s eyes (fixation point). Instructions with standardized guidelines were given to the subjects in advance, and before starting the measurements, the instructions were repeated in a short video (4 seconds of duration). To ensure that the participants learned the instructions correctly, a practice session was performed with 100 trials and the same amount of right and left targets. During the practice, the task was the same, only the probability of validly cued targets was modified (80%) to reinforce learning the cue. Subjects were reminded about fixating on the small fish’s eyes at the beginning of the task and in every break between blocks.

The visuospatial covert attention task was programmed and controlled using Presentation software (Neurobehavioral Systems, Albany, CA) and consisted of four blocks of 100 trials each. Each block contained 75 cued targets and 25 uncued ones. Between blocks, subjects could rest and after some minutes they decided when to start the following block. Left and right cues had the same probability of occurrence. The total duration of the task was approximately 20 minutes.

### MEG recordings and analysis

The brain activity related to the visuospatial covert attention paradigm was recorded using an Elekta Neuromag Triux system (Elekta Neuromag, Helsinki, Finland) for the subjects living in Central Finland. The recordings for the subjects from Southern Finland were performed in Aalto Brain Centre’s MEG core using Elekta Neuromag™ (Elekta Neuromag, Helsinki, Finland). The MEG devices in the two institutes are essentially the same. During the subject’s preparation, five head-position indicator coils were attached to their heads (three in the front and two behind both ears). Coil locations were determined using a 3-D digitizer in reference to three anatomical landmarks (nasion and pre-auricular points). At the beginning of the recording, their position with respect to the helmet was measured and continuously tracked during the whole measurement (Uutela et al., 2001) to correct for head movement during the analysis stage. With the use of the electrooculogram (EOG), vertical (two electrodes located at the external canthi of both eyes) and horizontal (two electrodes above and below the right eye) eye movements were monitored. During the measurements, the subjects were seated comfortably inside a magnetically shielded room and were instructed to avoid movements of the head and the eyes. Two response pads (left and right) were positioned on a table attached to the chair and located over the subject’s legs. The task was projected on a panel located 1 meter from the subject’s eyes.

The MEG signals were band-pass filtered at 0.03–330 Hz and sampled at 1000 Hz. The raw data were pre-processed using Maxfilter™ 2.2 software (Elekta Neuromag, Helsinki, Finland). A signal space separation method (SSS) (Taulu et al., 2004), removing external interference emerging during the measurement, was used in most of the subject’s recordings. SSS was replaced by the spatiotemporal signal space separation method (tSSS) (Taulu and Simola, 2006) (also included in Maxfilter 2.2 software) for the analysis of data from participants wearing braces or other internal magnetic sources. Head movements occurring during the measurement were also corrected by using Maxfilter 2.2 software. The rest of the analysis was performed with MNE-Python software (Gramfort et al., 2014). The analysis was performed with the brain activity recorded from the gradiometers. The time-frequency representations (TFRs) were calculated in a frequency range of 2–30 Hz and for a time period ranging from –0.2 to 1.4 seconds in reference to the onset of the cue. The frequency resolution used was 2 Hz. A Morlet wavelet transformation (Morlet et al., 1982) was used for this purpose with the number of cycles equal to half of each frequency value. For each channel, a modulation index (MI) was calculated with the following equation:

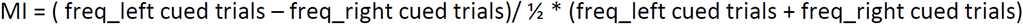

The resulting MIs were used to evaluate the visuospatial covert attention task-based modulation in the alpha band (8–12 Hz). Whole head TFRs were averaged together for the whole sample of 59 participants. Averages of three MEG channels in occipital regions where alpha increased (left) and alpha decreased (right) could be clearly identified were selected. The location of the left (contralateral) and right (ipsilateral) selections of MEG channels for the occipital region used for further analysis is shown in Fig. 3B. The difference between the alpha MI from the left minus right hemispheres was used to calculate the interhemispheric asymmetry of alpha MI during the task. Alpha MI in the left and right hemispheres and interhemispheric asymmetry were calculated for three different time-windows (0–400 ms, 401–800 ms, 801–1,200 ms).

MEG and behavioral data were analyzed in a blinded manner towards the group assignment to obtain an unbiased assessment of neurophysiological or cognitive outcomes.

### Attention and inhibitory control behavioral tasks

The participant’s speed of attention was assessed with the reaction time (RTI) test, which is a subtest in the Cambridge Neuropsychological Test Automated Battery (CANTAB) (Cambridge Cognition Ltd, 2006). The RTI test measures RT specifically and the speed of response towards an unpredictable target. During the unpredictable condition, a yellow spot appeared in any of five circles on the screen. Participants were instructed to retain their answers until they saw the yellow spot. Only then should they have touched, as fast as possible, the correct circle in the screen where the target had appeared. Subjects performed rehearsal trials until they understood the task and, afterward, 15 task trials. Scores were based on their RT (ms) and in movement time (ms).

The participants’ inhibitory control skills were assessed with a modified Eriksen Flanker task (Eriksen and Eriksen, 1974). In each trial, the target (central arrow) was flanked by non-target stimuli (surrounding arrows). During the compatible condition, the participants needed to report as fast as possible the target’s direction (left or right). During the incompatible condition, the subjects needed to report as fast as possible the opposite direction of the target (left button for the right direction of the target and right button for the left direction of the target). In both conditions, congruent trials (the target pointing in the same direction as the non-targets) and incongruent trials (the target pointing in the opposite direction to the non-targets) were included with the same probability of appearance. Accuracy (in percentages) and RT (in ms) were recorded for congruent and incongruent trials separately from each condition.

### Behavioral performance analysis

During the MEG measurement, behavioral responses from the visuospatial covert attention task were collected and analyzed. Incorrect responses (less than 2.5% of the data) and reaction times below or above 2.5 standard deviations from the mean in each condition for each participant (less than 2.5% of the data) were considered as outliers and excluded from further analysis. Reaction times under 250 ms were considered too short to reflect selective attention processing and were removed from the analysis. There was no difference in the number of cued and uncued targets from the visuospatial covert attention task used for further analysis for the whole sample divided by sex, the two groups based on AF and for the two groups based on PA.

Average reaction times for cued and uncued targets were calculated separately for each subject. A cueing effect index for RT was calculated as the difference between the reaction times for uncued minus cued targets. Accuracy was calculated as the percentage of correct responses. The cueing effect index for accuracy was calculated as the difference between the number of correct responses for cued minus uncued targets. Accordingly, average reaction times, accuracy for cued and uncued targets and the cueing effect indexes were used in further analysis.

In addition to typical task performance measures (accuracy and RT), the EZ model was used to analyze the results from the visuospatial covert attention task. Three new variables (drift rate, boundary separation and non-decision time) were calculated for each participant according to Wagenmakers et al. (2007). Drift rate, boundary separation, and non-decision time are determined from the mean RT (MRT), the variance of RT (VRT), and the proportion of correct responses (Pc). Thus, drift rate and boundary separation are calculated based on VRT and Pc, while MRT is used only to calculate non-decision time (for more details and R code see Wagenmakers et al. 2007).

### Statistical analysis

Statistical analysis was performed with IBM SPSS Statistics for Windows, Version 24.0 (Armonk, NY: IBM Corp.). Variables resulting from behavioral measures were assessed for normality. Only variables showing normal distribution were included in the statistical analysis. Also, the variables regarding accuracy from cognitive tests were removed from the final analysis due to reaching ceiling effects (more than 50% of the participants’ accuracy values were near the upper limit of the task range). Resulting variables used for further analysis per test were: (1) Visuospatial covert attention task (RT for cued and uncued targets, cueing effect for RT, drift rate, boundary separation and non-decision time); (2) Flanker task (RT for compatible conditions) and (3) RTI task (RT for five-choice condition, movement time for five-choice condition). All brain measures were included in future analyses.

We conducted two repeated measures analysis of variance (rANOVA) on the modulation index of alpha power, one for each between-subjects factor (PA and AF groups). Within-subjects factors were hemisphere (left, right) and time-windows (1^st^, 2^nd^, 3^rd^). Subsequent follow-up tests with rANOVA (separately for each time-window), were conducted for AF and PA groups as between-subjects factors to clarify the more detailed differences revealed by the primary analysis of variance tests.

Bivariate Pearson correlation coefficients were used to describe the associations between relevant brain data (interhemispheric asymmetry of alpha MI during the third time-window), cognitive measures (visuospatial covert attention task, Flanker task, and RTI test) and PA or AF variables (MVPA and minutes until exhaustion in the shuttle run test).

Because of the differences found in brain data for PA and AF groups we wanted to test the existence of a sex effect in the behavioral performance. Unfortunately, we could not do the same with the brain data because of the low signal to noise ratio, which could compromise the reliability of the potential results.

To test our hypothesis on whether oscillatory dynamics at the alpha band mediates the effect of PA or AF on cognition, a mediator analysis was performed by using MPlus software version 8.0 (Muthén and Muthén, 1998–2017). Before the analysis, we tested whether the assumptions for using the mediator model (normality and significant correlations between the three variables included in each model [independent variable, dependent variable, and moderator]) were met. Combinations with significant correlations of the mediator with independent and dependent variables were also included in the analysis. All possible models where alpha modulatory activity could mediate cognition improvements due to PA or AF levels were tested. The resultant combination was tested with a mediator model using MPlus software using a bootstrap of 1000 samples. Full information of maximum likelihood (FIML), which accounts for missing values at random (MAR) and includes all available data, was used.

In the mediation analysis, the bootstrap confidence interval was used, as it makes no assumption about the shape of the sampling distribution of the indirect effect (*ab*; Hayes and Rockwood, 2017). In bootstrapping, the indirect effect is estimated by randomly resampling cases from the dataset and estimating the model and resulting indirect effect in the bootstrap sample (Hayes and Rockwood, 2017; Preacher and Hayes, 2004). An empirical representation of the sampling distribution of the indirect effect is built by repeating this process 1,000 times. By using various percentiles of the bootstrap distribution, a confidence interval for the *ab* is constructed. Mediation is tested by determining whether the 95% confidence interval contains zero (Fritz and MacKinnon, 2007). If the interval is above or below zero, this supports mediation, and if the interval includes zero, it does not provide evidence of mediation. As recommended by Mackinnon (2008), the bias-corrected and accelerated bootstrap confidence intervals for the indirect effect were estimated using 1,000 bootstrap samples.

## ACKNOWLEDGMENTS

This study was supported by the Academy of Finland [grant numbers 273971, 274086 and 311877]. We would like to thank Ole Jensen for sharing the visuospatial covert attention task and his guidance in data analyses. For their valuable comments on this manuscript we would like to thank Ole Jensen, Georgia Gerike and Krista Lehtomäki. We also thank Janne Rajaniemi for his valuable help in the data collection. The authors declare no conflicts of interest.

